# Resting state fMRI detects pathological functional brain connectivity in the Ndufs4 knockout mouse model of Leigh syndrome during the pre-symptomatic stage of the disease

**DOI:** 10.64898/2026.10.22.760059

**Authors:** Chao Ciu-Gwok Guo, Melissa van de Wal, Samir el Bouazzati, Noortje M.P. Overwater, Lianne Remie, Werner Koopman, Joanes Grandjean, Judith R Homberg

## Abstract

Leigh syndrome (LS) is a progressive encephalopathy and generally regarded as the most common inborn mitochondrial disease. Mutations in mitochondrial respiratory complex I are common in patients with LS and a mutational hotspot has been identified in the gene that encodes the NDUFS4 subunit of respiratory complex I. The *Ndufs4* knockout mouse (*Ndufs4* KO) model of LS closely resembles the phenotypes of LS patients and show like the patient brain lesions when the symptoms are manifested. It is speculated but unproven that brain functional connectivity may precede the onset of symptoms. Therefore, the present study was set out to investigate the progression of brain functional connectives in *Ndufs4* KO mice before and after the manifestation of LS-like symptoms using *in vivo* resting-state functional magnetic resonance imaging (rs-fMRI). Since the brain lesions may be caused by increased reactive oxygen species (ROS) in the brain, we also examined the potential protective effect of KH176, a small molecular for modulating ROS levels, on brain functional connectivity. Rs-fMRI results showed that in *Ndufs4* KO mice the functional connectivity of the piriform cortex, motor cortex, barrel cortex, visual cortex, auditory cortex, prefrontal cortex, lateral striatum, ventral striatum cortex and amygdala was increased, starting from the pre-symptomatic stage (middle stage) to the symptomatic stage (late stage) of the disease. KH176 had no effect on the changes in functional connectivity in the mice. In conclusion, these results show that there are abnormalities in functional connectivity in Ndufs4 KO mice during the pre-symptomatic phase of the disease and that KH176 treatment does not modify the connectivity changes.

## Introduction

Mitochondria are cellular organelles that are present in virtually all eukaryotic cells and generate the vast majority of cellular energy in the form of adenosine triphosphate (ATP) via nutrient-dependent pathways. Mutations in nuclear DNA (nDNA) or mitochondrial DNA (mtDNA) encoded subunits of the oxidative phosphorylation system result in a heterogeneous group of multi-system diseases affecting 1:5,000 live births (Smeitink et al., 2001; Lightowlers et al., 2015; Gorman et al., 2016). Mutations in complex I (CI) encoding subunits induce isolated CI-deficiency, among which is Leigh syndrome (LS; OMIM #256000), also referred to as subacute necrotizing encephalopathy. Leigh syndrome is frequently caused by mutations in the *NDUFS4* (NADH:Ubiquinone Oxidoreductase Subunit S4) gene, which encodes an accessory subunit of CI that has been suggested to be involved in the assembly and/or stabilization of CI (Kahlhöfer et al., 2017; Adjobo-Hermans et al., 2020). All currently known pathogenic mutations in the *NDUFS4* gene result in an isolated CI-deficiency or pediatric LS. LS patients develop symptoms early in life, including progressive encephalopathy, failure to thrive, hypotonia, psychomotor retardation, breathing difficulties, recurrent vomiting, dysphagia, nystagmus, ataxia, neuropathy, hypertrophic cardiomyopathy, loss of vision, impaired hearing, seizures, lactic acidosis and eventually early death (Budde et al., 2003; Anderson et al., 2008; Assereto et al., 2014; Ortigoza-Escobar et al., 2016; Lamont et al., 2017). The manifestation of these symptoms in individuals with LS is often attributed to lesions in various brain structures (Budde et al., 2003; Anderson et al., 2008; Assouline et al., 2012; Assereto et al., 2014; Ortigoza-Escobar et al., 2016; Lamont et al., 2017). However, the contribution of these specific alterations in brain function preceding symptom manifestation remains unclear. Studies on other neurodegenerative diseases, such as Alzheimer’s disease (AD), have demonstrated impaired resting state functional connectivity (rs-fMRI) in individuals of AD before the emergence of symptoms (Gonneaud et al., 2021). Building on this insight, we hypothesize that alterations in brain functional connectivity may precede the onset of symptoms in LS patients. Unfortunately, there is currently a lack of available data to examine the functional connectivity of the brain in individuals before LS develops (Ardissone et al., 2021). A viable approach to address this knowledge gap involves longitudinal monitoring of brain functional connectivity in animal models of LS using rs-fMRI. This method allows for the continuous observation of changes in brain functional connectivity over time, offering valuable insights into the disease progression of LS. Such longitudinal studies may provide a deeper understanding of the temporal relationship between altered brain functional connectivity and the progression of LS symptoms.

The *Ndufs4* mouse model currently represents the best model to study LS (van de Wal et al., 2022). Whole body *Ndufs4* knockout (KO) mice appear healthy until postnatal day (PD) 35 and show similar behaviour as wildtype (WT) littermates. *Ndufs4* KO mice start to develop symptoms from PD35 onwards that closely mimic the human phenotype. Symptoms progressively worsen and mice die at ∼PD50 (Kruse et al., 2008). Aside from the overt clinical symptoms apparent in *Ndufs4* KO mice during the late disease stage (>PD39), noteworthy molecular changes manifest during the pre-symptomatic disease stage from PD26 (middle disease stage) onwards. Specifically, mice during the pre-symptomatic stage exhibit progressive microgliosis (Quintana et al., 2012; de Haas et al., 2017), reduced oxygen consumption in lesion-prone brain regions (Kayser et al., 2016) and alterations in glutamate/α-ketoglutarate metabolism (Johnson et al., 2020). In line with the disease progression observed in these studies, we systematically assessed brain functional connectivities of the mice at three distinct stages: the early pre-symptomatic stage (PD20-23), the middle pre-symptomatic stage (around PD26-29), and the late symptomatic stage (PD 39-40) using rs-fMRI.

Increased oxidative stress has been observed in CI-deficient cells models (Verkaart et al., 2007; Valsecchi et al., 2013) and in the retina (Song et al., 2017) and brain regions of *Ndufs4* KO mice (Quintana et al., 2010). In response to inflammation and oxidative stress, cells other than adipocytes can form lipid droplets (LDs). LD accumulation was observed in glial cells of lesion-prone brain regions of *Ndufs4* KO mice at PD23. LD accumulation by itself did not result in neurodegeneration but required ROS-induced lipid peroxidation. Reducing LD accumulation and lipid peroxidation in glial cells via targeted lipase overexpression and/or lowering reactive oxygen species (ROS) levels delayed the onset of neurodegeneration (Liu et al., 2015). In this sense, we hypothesized that modulating ROS levels may improve the phenotype of *Ndufs4* KO mice and specifically focussed on the effects of the ROS scavenger KH176 (sonlicromanol; (S)-6-hydroxy-2,5,7,8-tetramethyl-N-((R)-piperidin-3-yl)-chroman-2-carboxamide hydrochloride, (Patent WO2014011047 A1) on brain functional connectivity. KH176 has previously been proven to successfully modulate intracellular ROS levels (Beyrath et al., 2018)and improve several symptoms of *Ndufs4* KO mice (de Haas, 2017).

In this study, we assessed brain functional connectivity in *Ndufs4* WT and KO mice throughout the early pre-symptomatic, middle pre-symptomatic, and late symptomatic stages of the disease using rs-fMRI. Additionally, we extended this investigation to include mice from both genotypes that had undergone treatment with KH176 during the same three stages. We assessed functional connectivity within 17 canonical whole-brain networks, including networks involved in sensory processing (piriform cortex, auditory cortex, barrel field cortex, visual cortex), motor functions (dorsal striatum, ventral striatum, lateral striatum, motor cortex, supplementary motor cortex), integration of motor and sensory information (thalamus), emotion (amygdala, limbic cortex) and cognition (prefrontal cortex, dorsal hippocampus, ventral hippocampus). Our findings revealed that, in *Ndufs4* KO mice, nine networks, including piriform cortex, motor cortex, barrel field cortex, visual cortex, auditory cortex, prefrontal cortex, lateral striatum, ventral striatum, and amygdala showed increased functional connectivity starting from the middle stage compared to WT mice. Interestingly, no observable treatment effect of KH176 was noted in *Ndufs4* KO mice across all three disease stages. These results shed light on specific aberrations in functional connectivity in *Ndufs4* KO mice and suggest that KH176 treatment may not significantly impact these connectivity patterns throughout disease progression. Additionally, these results imply that rs-fMRI may help understanding the progression of LS symptoms.

## Materials and methods

### Animals and treatment

Initial breeding pairs of heterozygous *Ndufs4* mice (B6.129S4-Ndufs4 tm1.1Rpa/J background) were provided by Jackson Laboratory and were intercrossed to produce *Ndufs4* WT and KO in our breeding facility. Toe phalanx cuts were taken from pups for identification and for PCR genotyping at PD8-9. The primers for genotyping were 5’-GAT AGG GGA CTG GAC TAA CAG C-3’ and 5’-CAC TCC ACT GGA 63.2 °C GAG GAG C-3’. Mice were group-housed under controlled conditions (temperature: 20-22°C and humidity: 50-70%) with food (Ssniff VRF1, V1524-703, provider Bio Services, (irradiated 25 kGy)) and water provided *ad libitum*. Mice were maintained on a 12-hour light/dark cycle. Mice were weaned starting at PD20-21. Mice were housed in the presence of a wildtype littermate to provide warmth, and food was provided on the bottom of the cage. All mice were daily weighted and clinical symptoms were monitored. Mice were used for imaging and sacrificed at early-stage disease (PD20-23), middle-stage disease (PD26-29), and late-stage disease (PD39-40) as indicated in **Figure 1**. All animal experiments were conducted in accordance with national and local guidelines for animal experiments. The experiments were approved by the Central Committee Animal Experiments (The Hague, the Netherlands).

**Figure 1.**
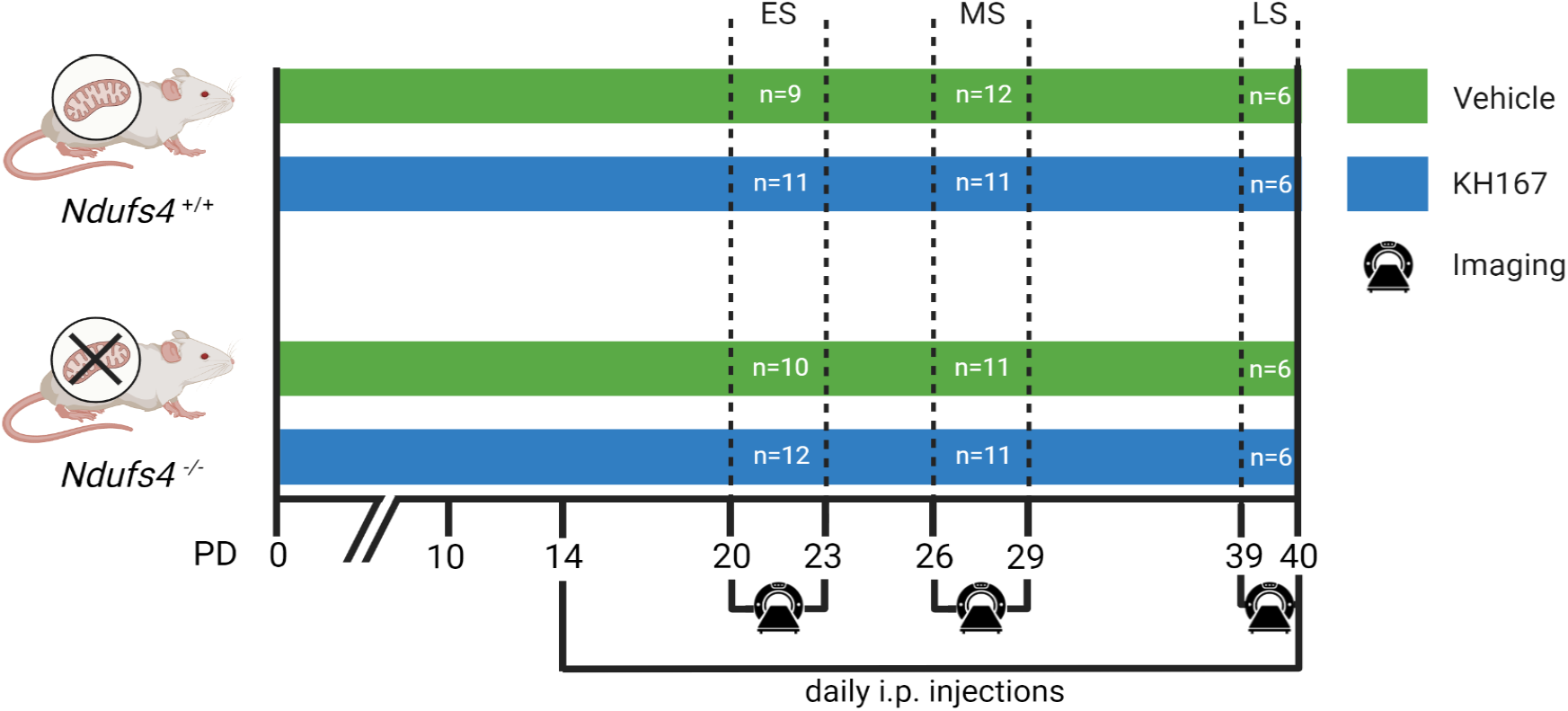
Overview of the experimental procedure. *Ndufs4* WT and *Ndufs4* KO mice were randomly assigned to one of the treatment groups (KH176 or the vehicle). The body weight of the mice was monitored from postnatal day (PD)10 until the day of imaging. Mice received daily injections of either vehicle (saline; early disease stage (ES) WT n=9, KO n=10; middle disease stage (MS) WT n=12, KO n=11; and late disease stage (LS) WT n=6, KO n=6) or KH176 (early disease stage WT n=11, KO n=12; middle disease stage WT n=11, KO n=11; and late disease stage WT n=6, KO n=6) from PD14 onwards. Mice were imaged on PD20-23 (early disease stage), PD26-29 (middle disease stage) or PD39-40 (late disease stage).

### Compound administration

Starting at PD14, mice were randomly assigned to receive either vehicle (saline) or KH176 ((S)-6-hydroxy-2,5-,7,8-tetramethyl-N((R)-piperidin-3-yl)chroman-2-carboxamide hydrochloride; Patent WO2014011047 A1, kindly provided by Khondrion B.V.). Both male and female mice were used and were equally divided among treatment groups to our best. KH176 was administered twice daily (10 mg/kg body weight) via intraperitoneal (I.P.) injections as previously described (de Haas et al., 2017).

### Data acquisition

MRI was performed using a Bruker BioSpec 117/16USR UHF small animal MR system (Bruker BioSpin MRI, Ettlingen, Germany) operating at 500 MHz (11.7 T) with an actively shielded gradient of 760 mT/m. The system was operated using Bruker Paravision 6.0.1 software. A circular quadrature volume coil (Bruker BioSpin AG, Fällanden, Switzerland) was used for signal transmission and a single-element receive-only coil (Bruker BioSpin AG, Fällanden, Switzerland) was used for signal reception. Low-dose isoflurane anaesthesia (Pharmachemie BV, Haarlem, the Netherlands) was supplied in a 20% O_2_/80% air mixture using a Tec-3 isoflurane vaporiser (Delta Medic, Ijsselstein, the Netherlands). Mice were exposed to isoflurane (1%-4%) during induction and lasted for about 1-3 min until breathing became visibly slower. The concentration of isoflurane was reduced to 0.75% during positioning in the MRI scanner and later reduced to 0.5% for maintenance throughout the imaging procedure. Subcutaneous injections of dexmedetomidine (Dexdomitor, Orion Phama, Vetoquinol, Espoo, Finland) in saline (0.5 mg/mL) at 0.15 mg/kg body weight were injected using 0.3L insulin syringes (30G x 8 mm needle). 20 minutes after the injection, rs-fMRI was performed. Ear bars were used to immobilize the head of mice during imaging. A rectal fiber optic probe thermometer (SA Instruments, Stony Brook, NY, USA) was used to measure the body temperature of the mice and temperature was maintained at 37 °C using warm water supplied by a Julabo U3 circulating water bath (Retsch BV, Aartselaar, Belgium). Respiration rate (breaths/min) and heart rate (beats/min) were monitored throughout the imaging procedure by using an air pressure sensitive sensor and MR-compatible small animal monitoring and gating system (SA Instruments, Stony Brook, NY, USA). After standard adjustments, coil tuning, coil matching, and shimming, a T2-weighted anatomical image was acquired. Imaging parameters for the anatomical scan are: TR=2500 ms; TE=30 ms; FOV=17x9 mm; matrix size=120x60x20; spatial resolution= 0.142x0.150x0.450 mm. Functional connectivity between brain regions was imaged using a rs-fMRI protocol with the following parameters: TR = 1000 ms; TE = 11.7 ms; FOV = 17 x 9 mm; matrix size = 60 x 30 x 20 x 720; spatial resolution = 0.283 x 0.300 x 0.450 mm; acquisition time =12 min. After the imaging procedure was finished, mice were sacrificed by decapitation.

### rs-fMRI analysis

Functional connectivity between brain regions, as measured by the rs-fMRI, was analyzed using RABIES version 0.4.5 (https://github.com/CoBrALab/RABIES). rs-fMRI data was transformed to BIDS format before analysis. Pre-processing steps included anatomical denoising, functional denoising, functional-anatomical alignment, distortion correction, head motion realignment and common space alignment of the T2-weighted anatomical image to the DSURQE template(Dorr et al., 2008). A manual quality control step was done on the aligned images to check for any misalignments. Dual regression was carried out using 17 components representing whole-brain networks previously identified (Zerbi et al., 2015), including the amygdala, dorsal and ventral hippocampus, dorsal, lateral and ventral striatum, thalamus, auditory cortex, barrel field cortex, cingulate cortex, limbic cortex, motor cortex, olfactory cortex, piriform cortex, prefrontal cortex, supplementary cortex and visual cortex (**Supplementary Figure S1**). We extracted the averaged connectivity within each network to carry out the statistical analysis (Grandjean et al., 2016).

### Statistical analysis

Data were analyzed using a mixed-effect linear regression model in R with lme4 package (Bates et al., 2015). For brain functional connectivity, disease stage, genotype, treatment, and interaction between of genotype*treatment and genotype*stage were assessed as fixed effects, and age and body weight on the day of imaging were assessed as the random effects in the model. For body-weight analysis, genotype, treatment, age, and interactions of genotype*age and genotype*treatment treatment*age were assessed as fixed effects, and sex and subject were assessed as random effects in the model. To accept the uncertainty and move beyond the p < 0.05 in statistical inference, the estimate (E) and 95% bootstrapped (5000 times) confidence interval (CI) of each fixed effect was provided (Wasserstein et al., 2019). If the range of 95% bootstrapped confidence interval did not include 0, the effect was deemed strong (Guo et al., 2021). If the fixed effect was strong, a following-up analysis was performed to test the effect in each stage of the disease. Figures were plotted with Python (version 3.8) or R (version 4.2).

### Data availability

The raw data in BIDS format is available under the terms of the CC0 license at doi:10.18112/openneuro.ds004976.v1.0.0

## Results

### Body weight

Body weights of all mice were daily measured starting from PD10 until the day of *in vivo* imaging. Growth curves between genotypes and treatment groups were compared (**Figure 2**). In general, KO mice had lower body weight than WT mice. KH176 treatment had no strong effect on body weight in both KO and WT mice. On the day of brain imaging, KO mice had lower body weight than WT mice in each disease stage (early, middle, and late stage). See **Table 1** for the estimation of genotype, treatment, genotype*treatment effects, and bootstrapped confidence intervals of each effect.

**Table 1.** Estimation of confidence intervals for each effect in across the ages and the date of brain imaging.

| Across the stage |  | On the day of imaging |  |  |
| --- | --- | --- | --- | --- |
| Fixed effect | All stages | Early pre-symptomatic disease stage | Middle pre-symptomatic disease stage | Late symptomatic disease stage |
| Genotype | E = -0.712, CI = [-0.892, -0.536] | E = -0.662, CI = [-1.0552, -0.2719] | E = -1.21, CI = [-1.572, -0.835] | E = -2.78, CI = [-3.169, -2.403] |
| Treatment | E = -0.0131, CI = (-0.168, 0.194] | E = 0.0446, CI = [-0.2604, 0.3554] | E = 0.107, CI = [-0.1883, 0.4071] | E = -0.109, CI = [-3.169, 0.2435] |
| Genotype:Treatment | E = 0.0222, CI = (-0.161, 0.200] | E = -0.246, CI = [-0.5394, 0.048] | E = -0.153, CI = [-0.4642, 0.14] | E = 0.203, CI = [-0.1317, 0.5491] |
| Age | E = 0.423, CI = [0.416, 0.429] |  |  |  |
| Genotype:Age | E = -0.0881, CI = (-0.0943, -0.0819] |  |  |  |

**Figure 2.**
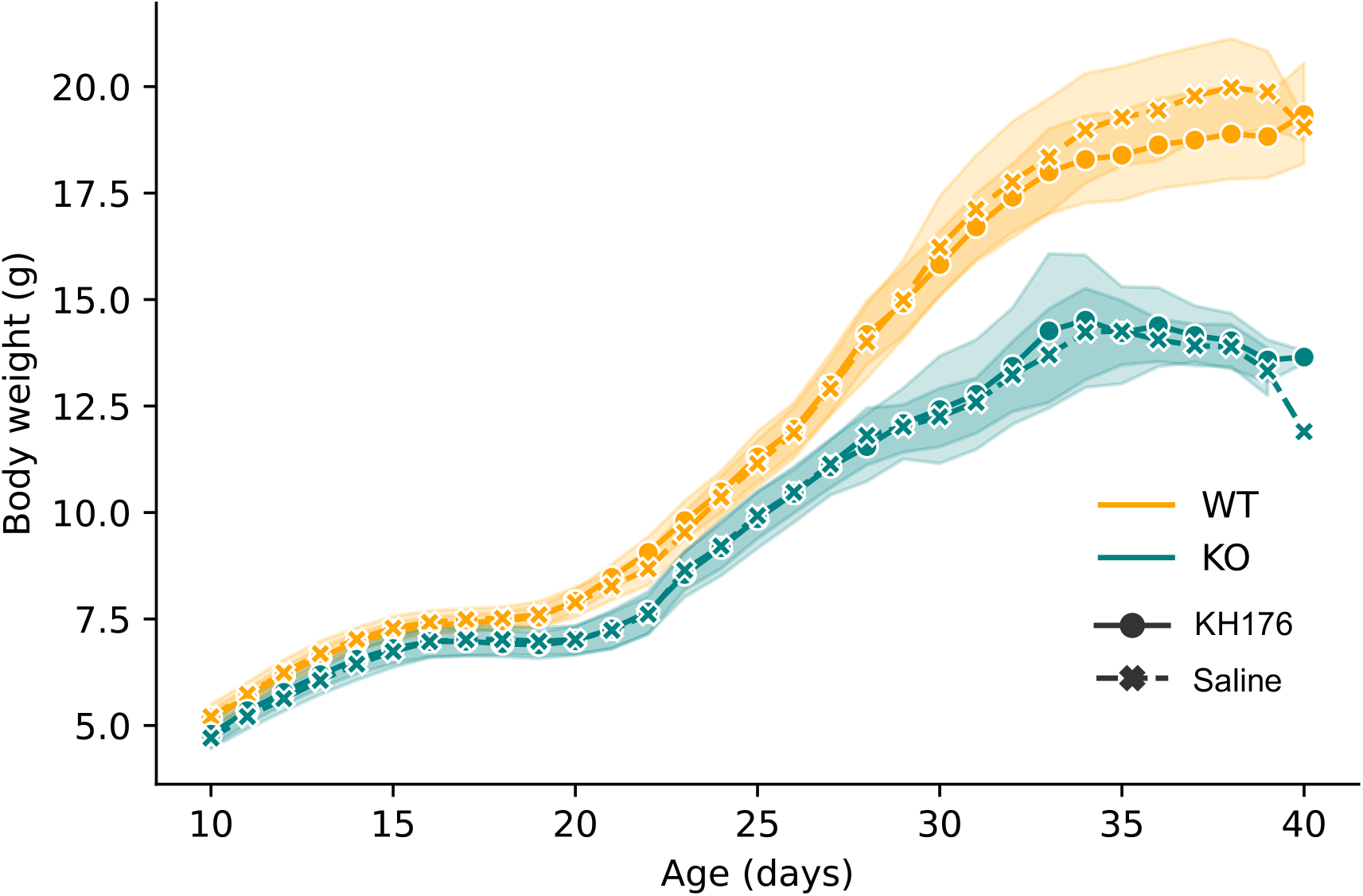
Body weights of *Ndufs4* WT (yellow lines) and *Ndufs4* KO (green lines) treated with vehicle (dashed line) or KH176 (solid line) mice from PD10 onwards. The solid and dash lines are the mean value of body weight. The transparent strips around the lines are the 95% confidence intervals.

### Functional connectivity

Functional connectivity within canonical networks of the mouse brain was assessed using rs-fMRI. Rs-fMRI data were analyzed using dual regression, and parameter estimates for 17 validated functional networks (cortical, limbic system, basal ganglia, thalamic) were obtained. All data are depicted in **Figure 3**. In general, we found that genotype and genotype*stage effects or one of them was strong in all brain networks. The treatment effect was not strong in all brain networks. Further analysis showed that no strong differences were observed between WT and KO mice in the early pre-symptomatic disease stage (PD20-23). Interestingly, strong differences in the piriform cortex network, motor cortex network, barrel field cortex 1, visual cortex network, prefrontal cortex network, lateral and ventral striatum network and amygdala functional network connectivities were found starting from the middle pre-symptomatic disease stage (PD26-29) of *Ndufs4* KO mice compared to WT littermates. All analyzed functional networks showed a strong difference between WT and KO mice in the late symptomatic disease stage (PD39-30). Strikingly, no strong treatment effects were observed for each brain region in each disease stage, indicating that KH176 treatment did not affect functional connectivity in our study. All genotype effects for each functional connectivity and disease stage are summarized in **Table 2**. Further details can be found in **Supplementary materials** of brain network estimations.

**Table 2:**
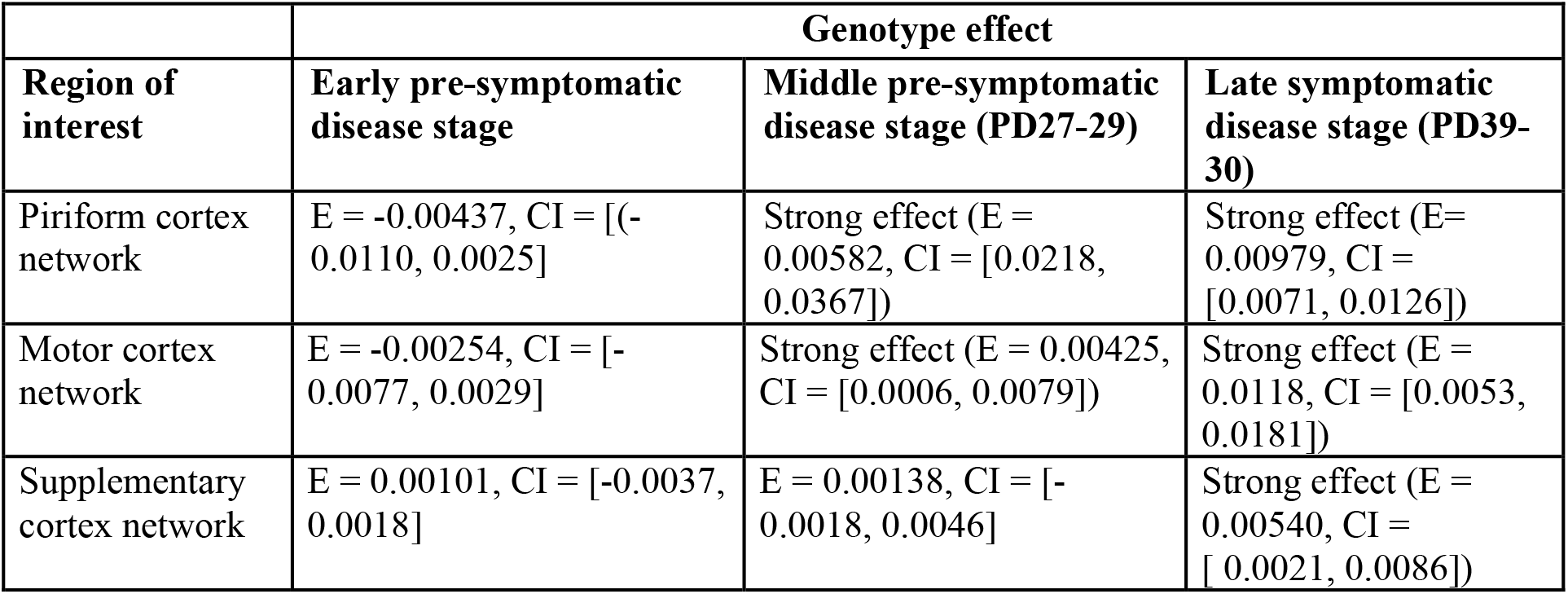
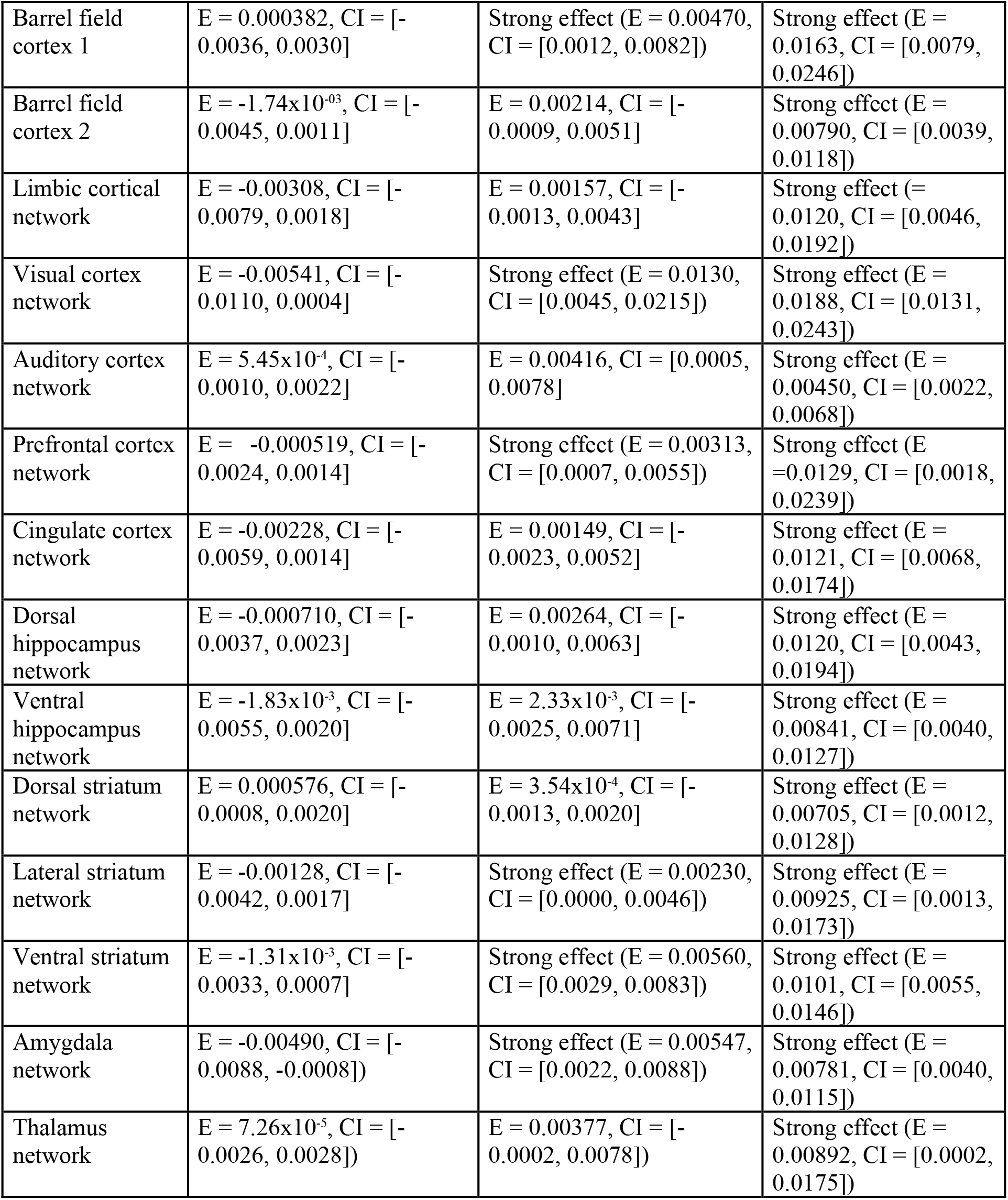
Genotype effect on functional connectivity in the different functional networks for each disease stage. Estimation (E) and bootstrapped confidence interval (CI) are given.

**Figure 3.**
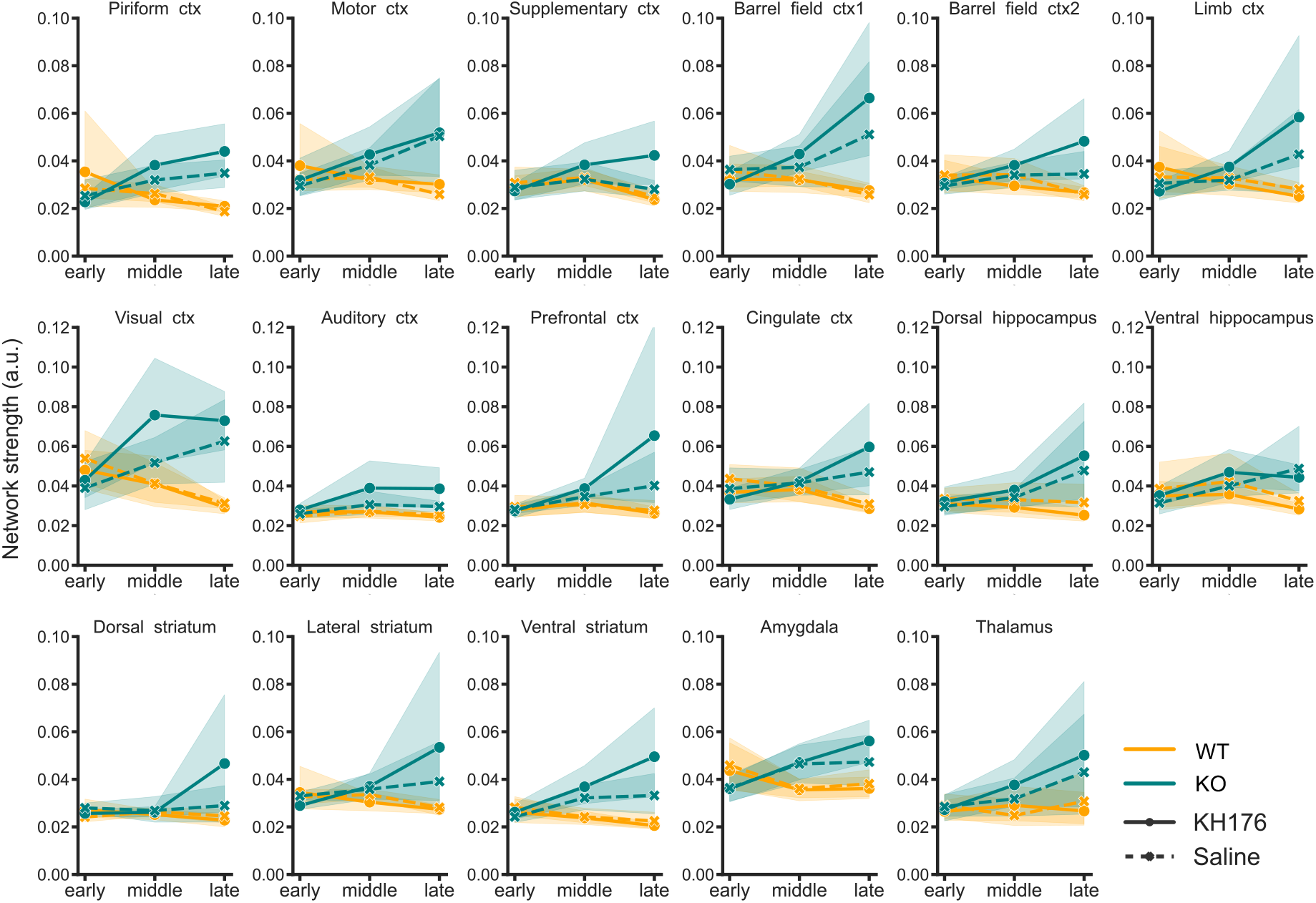
Functional connectivity in selected regions of interest (ROIs). Network strength was measured during three stages of LS disease progression in vehicle (solid lines) and KH176-treated (dashed lines) *Ndufs4* WT (yellow) and KO (green) mice (early pre-symptomatic stage (PD 20-22); middle pre-symptomatic stage (PD 26-29), and late symptomatic stage (PD 39-40)). Network strength is expressed in arbitrary units (a.u.). ctx is short for cortex. ROIs were drawn on a mean T2-weighted anatomical image of Ndufs4+/+ mice (see Supplementary Figure S3). The dots are the mean value of network strength, the transparent stripes are 95% of bootstrapped confidence intervals.

## Discussion

In this study, we aimed to investigate the effect of *Ndufs4* gene deletion on brain functional connectivity using rs-fMRI during the early (PD20-23) and middle (PD26-29) pre-symptomatic and late (PD39-40) symptomatic disease stages. In addition, we evaluated the effect of the ROS modulator KH176 on brain functional connectivity in these disease stages. We demonstrated that *Ndufs4* KO mice exhibited higher levels of functional connectivity strength compared to WT littermates, in the pre-symptomatic middle disease stage (PD26-29) and symptomatic late disease stage (PD39-40). One interpretation for increased connectivity may be the manifestation of a larger demand on the vascular system to provide oxygen to metabolically inefficient cells in the brain in *Ndufs4* KO mice. Alternatively, increased connectivity may relate to reduced signal transmission efficacy. In a study using chemogenetics to reduce excitatory transmission, Rocchi et al. found increased functional connectivity (Rocchi et al., 2022). Both models seem plausible to explain the results from this study, each suggesting different possible interventions.

Next to brain abnormalities, increased oxidative stress has been observed in CI-deficient cell models (Verkaart et al., 2007; Valsecchi et al., 2013) as well as retina of *Ndufs4* KO mice as early as PD20 (Song et al., 2017). KH176 has been proven to successfully module ROS levels within cells (Beyrath et al., 2018) and to normalized lipid peroxidation, maintain microstructural coherence, reduce ganglion cell degeneration, and to improve motor performance and gait impairment in *Ndufs4* KO mice (de Haas, 2017). To this end, we hypothesized that KH176 would normalize the aberrant functional connectivity in *Ndufs4* KO mice. Surprisingly, KH176 was unable to restore the brain functional connectivity in *Ndufs4* KO mice in any of the analysed phases. Probably this is due to the small sample size in the late stage of the disease. This suggests that oxidative stress does not play a role in maintaining brain functional connectivity.

The specific network alterations that we observe may actually recapitulate pathological events linked to the pathology. For instance, bilateral spongiform lesions in *Ndufs4* KO mice have been observed in the vestibular nuclei (located in medulla and pons of hindbrain) and developed neurodegeneration in the olfactory bulb, optic chiasm, optic tract, superior colliculus, interpenduncular nucleus, lateral lemniscus, trapezoid body, cochlear nuclei, fastigial nucleus, inferior olivary complex, nodulus (X) and the uvula (IX) granular and molecular layer (Quintana et al., 2012; Liu et al., 2015; de Haas et al., 2016). However, these areas are not part of the canonical networks (Zerbi et al., 2015) observed in the mouse brain and are not amenable for analysis due to the limited resolution of rs-fMRI. The effects in these structures may be found in the higher order networks investigated. Neurological abnormalities are commonly observed in the OB of *Ndufs4* KO mice (Quintana, 2012). The motor cortex generates signals to direct movements of the body (Wise and Donoghue, 1986) and lesions in this region result in motor impairments (Heindorf et al., 2018). *Ndufs4* KO mice lose their motor function from PD30 onwards (Kruse, 2008; de Haas, 2016) and increased functional connectivity strength as already observed in the presymptomatic middle disease stage may contribute to this. The barrel field cortex is a region in the somatosensory cortex that is involved in processing spatial and textural information of the immediate surroundings (Petersen, 2019). The visual and auditory cortex are structures located in the cerebral cortex and are involved in processing visual and auditory information. *Ndufs4* KO mice develop blindness and deafness (Kruse, 2008) and increased functional connectivity in these brain regions in the middle presymptomatic disease stage indicate that alterations already occur during the presymptomatic phase. The prefrontal cortex is involved in cognition, motivation, reward and emotion (Dalley et al., 2004) and the amygdala has been associated with emotional regulation, such as fear and anxiety (LeDoux, 2003).

The striatum consists of different subdivision (ventral, dorsal, lateral), which are connected with different cortical and subcortical structures and form channels that process information related to limbic, associative, and sensorimotor functions (Burton et al., 2015). For example, the ventral striatum has been suggested to serve as a gateway where information from the amygdala and prefrontal regions are processed and can influence motor output to guide behavior (Roesch et al., 2009). Increased functional connectivity in the striatum may indicate reduced anatomical connectivity and loss of diversity in interaction with the amygdala and prefrontal cortex. Collectively, our data suggests that increased functional connectivity in the described brain regions contribute to disease progression of *Ndufs4* KO mice. Furthermore, we show that alterations can already be detected using rs-fMRI as early as the middle presymptomatic disease stage, before symptoms become apparent. In addition to functional connectivity analysis, various computational methods such as voxel-based morphometry, atlas-based analysis, and deep learning are extensively employed to characterize neurological abnormalities in different diseases and could give more insight in the pathophysiology of LS.

## Acknowledgment

We thank the animal caretakers of the central animal facility of the Radboud University Medical Center for their assistance with KH176 treatment, and Msc. Anthea Grum for her assistance with data collection. C.G. arranged and coordinated the schedule and timeline of the experiments. C.G., S.B. and L.R. executed the experiment. C.G., S.B., L.R. and M.V contributed to the manuscripts. J.G. provided valuable guidance and help for rs-fMRI data acquisition and image analysis. J.H, J.G. W.K. provided critical feedback on manuscript versions. S.B. and L.R were supervised by C.G.. C.G. and M.V. were supervised by J.G., J.H. and W.K.. This work is financially supported by Stichting Zeldzame Ziekten Fonds.

