## Supplementary materials for "Resting state fMRI detects pathological functional brain connectivity in the Ndufs4 knockout mouse model of Leigh syndrome during the pre-symptomatic stage of the disease"

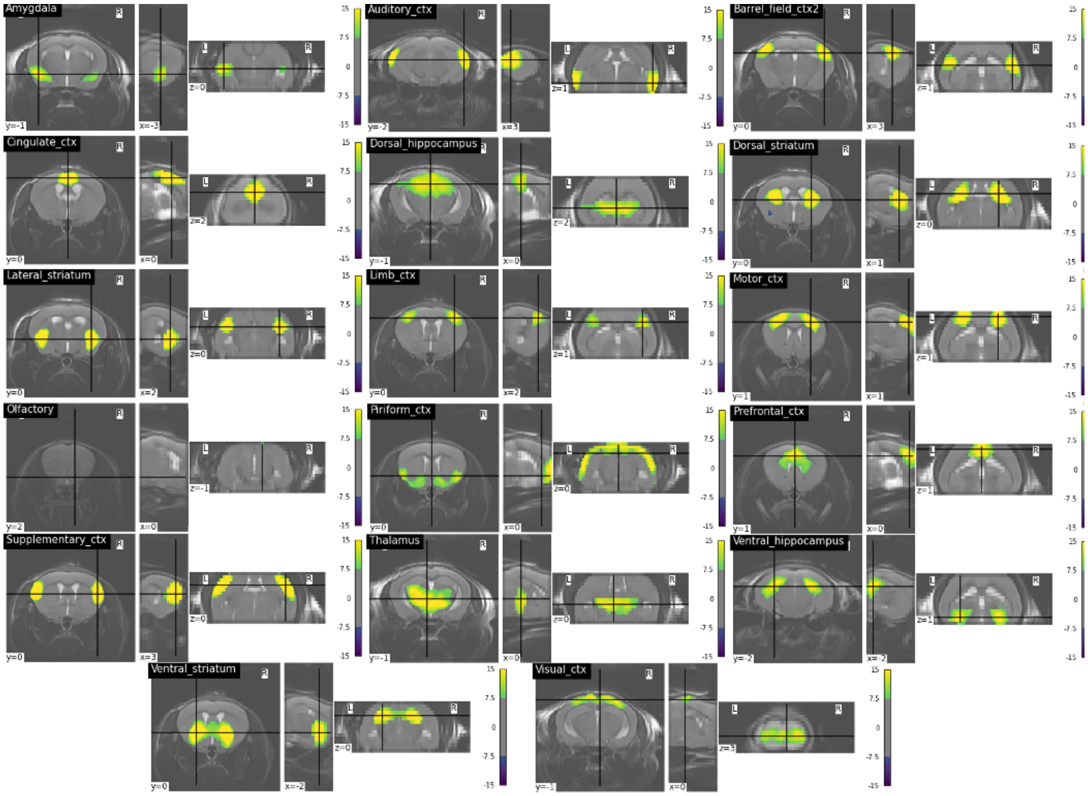


**Supplementary Figure S1.** Network masks overlaid on mean T2-weighted anatomical image of Ndufs4+/+ mice following RABIES preprocessing. The networks include the amygdala, auditory cortex, barrel field cortex, visual cortex, cingulate cortex, dorsal hippocampus, ventral hippocampus, dorsal striatum, ventral striatum, lateral striatum, limbic cortex, motor cortex, piriform cortex, prefrontal cortex, supplementary motor cortex, thalamus.

**Brain network estimations**

In the piriform cortex network, effects of genotype (E = 0.0041540, CI = [0.0015, 0.0067]) and the interaction between genotype and stage (genotype*stage) (E = 0.0045490, CI = [0.0019, 0.0073]) were strong. The effects of treatment (E = 0.0002557, CI = [-0.0023, 0.0029]), stage (E = 0.0006355, CI = [-0.0036, 0.0050]), genotype*treatment (E = -0.0002340, CI = [-0.0027, 0.0024]) were not strong. Follow-up analysis showed that the genotype effect was strong in the middle stage (E = 0.0058203, CI = [0.0218, 0.0367]) and late stage (E = 0.009793, CI = [0.0071, 0.0126]), but not in the early stage (E = -0.004369, CI = [(-0.0110, 0.0025]) of the disease.

In the motor cortex network, genotype (E = 0.0038551, CI = [0.0010, 0.0068]) and genotype*stage (E = 0.0055899, CI = [ 0.0028, 0.0084]) effects were strong. However, the effects of treatment (E = 0.0013664, CI = [-0.0014, 0.0042]), stage (E = 0.0030135, CI = [-0.0008, 0.0069]), and genotype*treatment (E = 0.0002341, CI = [-0.0026, 0.0031]) were not strong. Follow-up analysis showed that the genotype effect was strong in the middle (E = 0.0042489, CI = [ 0.0006, 0.0079]) and late stages (E = 0.0117528, CI = [0.0053, 0.0181]), but not in the early stage (E = -0.0025396, CI = [-0.0077, 0.0029]) of the disease.

In the supplementary cortex network, there were no strong effects of genotype (E = 0.0016604, CI = [-0.0002, 0.0035]), stage (E = 0.0006566, CI = [-0.0020, 0.0034]), treatment (E = 0.0007991, CI = [-0.0010, 0.0026]) or treatment*genotype (E = 0.0016499, CI = [-0.0002, 0.0035]), but there was a strong genotype*stage effect (E = 0.0026965, CI = [0.0009, 0.0045]). Follow-up analysis showed there were no strong genotype effects in both early (E = 0.0010071, CI = [-0.0037, 0.0018]) and middle (E = 0.001383, CI = [-0.0018, 0.0046]) stages, but a strong genotype effect in late stage (E = 0.005401, CI = [ 0.0021, 0.0086]). Since there was no strong genotype effect in the full model, the genotype effect in the late stage should be moderate.

In the barrel field cortex 1 network, the effects of genotype (E = 0.0057947, CI = [ 0.0029, 0.0087]), stage (E = 0.0037850, CI = [0.0001, 0.0075]) and genotype*stage (E = 0.0063727, CI = [0.0036, 0.0092]) were strong. However, the effect of treatment (E = 0.0005361, CI = [-0.0022, 0.0034]) and treatment*genotype (E = 0.0010789, CI = [-0.0018, 0.0039]) were not strong. Follow-up analysis showed that the genotype effect was strong in the middle stage (E = 0.004702, CI = [0.0012, 0.0082]) and late stage (E = 0.016296, CI = [0.0079, 0.0246]), but not in early stage (E = 0.0003819, CI = [-0.0036, 0.0030]) of the disease.

In the barrel field cortex 2 network, the effects of genotype (E = 0.00230788, CI = [0.0005, 0.0042]) and genotype*stage (E = 0.0037987, CI = [0.0020, 0.0056]) were strong, but the effects of stage (E = 0.0011575, CI = [-0.0014, 0.0037]), treatment (E = 0.0007114, CI = [-0.0011, 0.0026]), and treatment*genotype (E = 0.0017903, CI = [-0.0001, 0.0037]) were not strong. Follow-up analysis showed that the genotype effect was strong in the late stage (E = 0.007898, CI = [0.0039, 0.0118]), but not in the early (E = $-1.737\times10^{-03}$, CI = [-0.0045, 0.0011]) or middle stage (E = 0.0021367, CI = [-0.0009, 0.0051]) of the disease.

In the limbic cortical network, the effect of genotype*stage (E = 0.0058753, CI = [0.0031, 0.0086]) was strong, but the other effect were not strong (genotype: E = 0.0025413, CI = [-0.0003, 0.0054]; stage: E = 0.0023008, CI = [-0.0005, 0.0051]; treatment: E = 0.0009109, CI = [-0.0018, 0.0037]; genotype*treatment: E = 0.0012577, CI = [-0.0015, 0.0041]). Follow-up analysis showed that the genotype effect was strong in the late stage (E = 0.011973, CI = [0.0046, 0.0192]) but not in the middle (E = 0.0015660, CI = [-0.0013, 0.0043]) or early stage (E = -0.0030833, CI = [-0.0079, 0.0018]) of the disease.

In the visual cortex network, the effects of genotype (E = 0.008077, CI = [0.0040, 0.0123]), genotype*treatment (E = 0.004302, CI = [0.0002, 0.0084]), and genotype*stage (E = 0.009926, CI = [0.0059, 0.0140]) were strong, but not the treatment (E = 0.002434, CI = [-0.0016, 0.0064]) and stage (E = 0.002100, CI = [-0.0050, 0.0095]) effects. Follow-up analysis showed that the genotype effect was strong in the middle (E = 0.013009, CI = [0.0045, 0.0215]) and late (E = 0.018749, CI = [0.0131, 0.0243]) stages, but not in the early stage (E = -0.0054121, CI = [-0.0110, 0.0004]) of the disease. The treatment effect in each stage was not strong (early: E = -0.0005355, CI = [-0.0065, 0.0054], middle: E = 0.005306, CI = [-0.0029, 0.0132], late: E = 0.002087, CI = [-0.0036, 0.0078]).

In the auditory cortex network, the effects of genotype (E = 0.0029578, CI = [0.0013, 0.0047]) and genotype*stage (E = 0.0018543, CI = [0.0003, 0.0035]) were strong, but not the effects of stage (E = 0.0013874, CI = [-0.0010, 0.0038]), treatment (E = 0.0014811, CI = [-0.0001, 0.0031]), and genotype*treatment (E = 0.0014582, CI = [-0.0002, 0.0031]). Follow-up analysis showed that the genotype effect in the early (E = $5.449 \times 10^{-4}$, CI = [-0.0010, 0.0022]) or middle (E = 0.004156, CI = [0.0005, 0.0078]) stage was not strong, while it was strong in the late stage (E = 0.004503, CI = [0.0022, 0.0068]) of the disease.

In the prefrontal cortex network, the effects of genotype (E = 0.0035276, CI = [0.0006, 0.0065]) and genotype*stage (E = 0.0014582, CI = [(-0.0002, 0.0031]) were strong, but not the effects of stage (E = 0.0034560, CI = [-0.0001, 0.0068]), treatment (E = 0.0004903, CI = [-0.0025, 0.0035]), and genotype*treatment (E = 0.0011355, CI = [-0.0017, 0.0041]). Follow-up analysis showed that the genotype effect in the middle (E = 0.0031301, CI = [0.0007, 0.0055]) and late (E =0.012933, CI = [0.0018, 0.0239]) stages were strong, but not in the early stage (E = -0.0005193, CI = [-0.0024, 0.0014]).

In the cingulate cortex network, the effects of genotype (E = 0.0028745, CI = [0.0005, 0.0053]) and genotype*stage (E = 0.0054453, CI = [0.0030, 0.0079]) were strong, but not the effect of stage (E = 0.0012342, CI = [-0.0014, 0.0039]), treatment (E = -0.0007341, CI = [-0.0031, 0.0016]), or genotype*treatment (E = 0.0009668, CI = [-0.0015, 0.0034]). Follow-up analysis showed that the genotype effect in the late stage was strong (E = 0.012126, CI = [0.0068, 0.0174]), but not in the middle (E = 0.0014935, CI = [-0.0023, 0.0052]) or early (E = -0.0022774, CI = [-0.0059, 0.0014]) stage of the disease.

In the dorsal hippocampus network, the effects of genotype (E = 0.0032160, CI = [0.0006, 0.0058]) and genotype*stage (E = 0.0042446, CI = [0.0016, 0.0069]) were strong, but not the effects of stage (E = 0.0026005, CI = [(-0.0004, 0.0057]), treatment (E = -0.0008203, CI = [-0.0034, 0.0017]), and genotype*treatment (E = 0.0017355, CI = [-0.0007, 0.0043]). Follow-up analysis showed that the genotype effect in the late (E = 0.0119570, CI = [0.0043, 0.0194]) stage was strong, but not in the middle (E = 0.0026366, CI = [-0.0010, 0.0063]) or early (E = -0.0007096, CI = [-0.0037, 0.0023]) stage.

In the ventral hippocampus network, the effects of genotype (E = 0.0024627, CI = [0.0000, 0.0051]) and genotype*stage (E = 0.0040088, CI = [0.0015, 0.0065]) were strong, but not the effects of stage (E = 0.0015164, CI = [-0.0019, 0.0049]), treatment (E = -0.0005198 , CI = [-0.0030, 0.0020]), and genotype*treatment (E = 0.0021190, CI = [-0.0004, 0.0046]). Follow-up analysis showed that the genotype effect in the late (E = 0.0084072, CI = [0.0040, 0.0127]) stage was strong, but not in the middle (E = $2.330 \times 10^{-3}$, CI = [-0.0025, 0.0071]) and early (E = $-1.826 \times 10^{-3}$, CI = [-0.0055, 0.0020]) stage of the disease.

In the dorsal striatum network, the effects of genotype (E = 0.0019680, CI = [0.0003, 0.0037]) was strong, but not the effects of stage (E = 0.0009586, CI = [-0.0012, 0.0031]), treatment (E = 0.0002952, CI = [-0.0014, 0.0020]), genotype*stage (E = 0.0017038, CI = [-0.0001, 0.0035]), and genotype*treatment (E = 0.0002036, CI = [-0.0014, 0.0019]). Follow-up analysis showed that the genotype effect in the late (E = 0.007053, CI = [0.0012, 0.0128]) stage was strong, but not in the middle (E = $3.543 \times 10^{-4}$, CI = [-0.0013, 0.0020]) and early (E = 0.0005757, CI = [-0.0008, 0.0020]) stage of the disease.

In the lateral striatum network, the effects of genotype (E = 0.0030164, CI = [0.0006, 0.0055]) and genotype*stage (E = 0.0034232, CI = [0.0008, 0.0060]) were strong, but not the effects of stage (E = 0.0013295, CI = [-0.0016, 0.0043]), treatment (E = -0.0001544, CI = [-0.0026, 0.0023]), and genotype*treatment (E = 0.0002773, CI = [-0.0021, 0.0028]). Follow-up analysis showed that the genotype effect in the middle (E = 0.0023031, CI = [0.0000, 0.0046]) and late (E = 0.009245, CI = [0.0013, 0.0173]) stages were strong, but not in the early stage (E = -0.0012781, CI = [-0.0042, 0.0017]) of the disease.

In the ventral striatum network, the effects of genotype (E = 0.0040477, CI = [0.0024, 0.0056]) and genotype*stage (E = 0.0038016, CI = [0.0022, 0.0055]) were strong, but not the effects of stage (E = 0.0014677, CI = [-0.0015, 0.0046]), treatment (E = 0.0007814, CI = [-0.0008, 0.0024]), and genotype*treatment (E = 0.0013512, CI = [-0.0001, 0.0029]). Follow-up analysis showed that the genotype effect in the middle (E = 0.0056032, CI = [0.0029, 0.0083]) and late (E = 0.010116, CI = [0.0055, 0.0146]) stages were strong, but not in the early stage (E = $-1.308 \times 10^{-3}$, CI = [-0.0033, 0.0007]) of the disease.

In the amygdala network, the effects of genotype (E = $2.494 \times 10^{-3}$, CI = [0.0003, 0.0047]) and genotype*stage (E = $4.657 \times 10^{-3}$, CI = [0.0025, 0.0068]) were strong, but not the effects of stage (E = $1.695 \times 10^{-3}$, CI = [-0.0017, 0.0053]), treatment (E = $8.072 \times 10^{-5}$, CI = [-0.0021, 0.0022]), and genotype*treatment (E = $9.384 \times 10^{-4}$, CI = [-0.0012, 0.0031]). Follow-up analysis showed that the genotype effect in the middle (E = 0.0054734, CI = [0.0022, 0.0088]) and late (E = 0.007809, CI = [0.0040, 0.0115]) stages were strong, but not in the early stage (E = -0.0049024, CI = [-0.0088, -0.0008]) of the disease.

In the thalamus network, the effect of genotype (E = 0.0040141, CI = [0.0013, 0.0067]) was strong, but not the effects of stage (E = 0.0029915, CI = [-0.0001, 0.0061]), treatment (E = 0.0001513, CI = [-0.0029, 0.0026]), genotype*stage (E = 0.0013897, CI = [-0.0015, 0.0043]), and genotype*treatment (E = 0.0009451, CI = [-0.0016, 0.0037]). Follow-up analysis showed that the genotype effect in the late (E = 0.0089158, CI = [0.0002, 0.0175]) stage was strong, but not in the middle (E = 0.0037699, CI = [-0.0002, 0.0078]) and early (E = $7.255 \times 10^{-5}$, CI = [-0.0026, 0.0028]) stage of the disease.
